# Zinc Alpha-2-Glycoprotein (ZAG/AZGP1) secreted by triple-negative breast cancer promotes tumor microenvironment fibrosis

**DOI:** 10.1101/2024.03.04.583349

**Authors:** Surbhi Verma, Stephanie Dudics Giagnocavo, Meghan C. Curtin, Menusha Arumugam, Sandra M. Osburn-Staker, Guoying Wang, Aaron Atkinson, David A. Nix, David H. Lum, James E. Cox, Keren I. Hilgendorf

**Affiliations:** Department of Biochemistry, University of Utah School of Medicine, Salt Lake City, UT 84112, USA; Metabolomics, Proteomics and Mass Spectrometry Core, School of Medicine, University of Utah, Salt Lake City, UT, USA; Huntsman Cancer Institute, University of Utah, Salt Lake City, UT, USA

**Keywords:** ZAG, AZGP1, Zinc Alpha-2-Glycoprotein, cancer-associated fibroblasts, triple-negative breast cancer, secretome, adipocyte stem and progenitor cells, adipose tissue, tumor microenvironment, fibrosis

## Abstract

**Obesity is a predisposition factor for breast cancer, suggesting a localized, reciprocal interaction between breast cancer cells and the surrounding mammary white adipose tissue. To investigate how breast cancer cells alter the composition and function of adipose tissue, we screened the secretomes of ten human breast cancer cell lines for the ability to modulate the differentiation of adipocyte stem and progenitor cells (ASPC). The screen identified a key adipogenic modulator, Zinc Alpha-2-Glycoprotein (ZAG/AZGP1), secreted by triple-negative breast cancer (TNBC) cells. TNBC-secreted ZAG inhibits adipogenesis and instead induces the expression of fibrotic genes. Accordingly, depletion of ZAG in TNBC cells attenuates fibrosis in white adipose tissue and inhibits tumor growth. Further, high expression of ZAG in TNBC patients, but not other clinical subtypes of breast cancer, is linked to poor prognosis. Our findings suggest a role of TNBC-secreted ZAG in promoting the transdifferentiation of ASPCs into cancer-associated fibroblasts to support tumorigenesis.**

## Introduction

Approximately 1 in 8 women will develop breast cancer in their lifetime, and more than 40,000 women in the United States die from breast cancer every year (Vegunta et al., 2022). Breast cancer is clinically subdivided into subtypes based on the expression of three cellular markers, estrogen receptor, progesterone receptor, and human epidermal growth factor receptor 2. Triple-negative breast cancer (TNBC) does not express any of these markers, accounts for 10-20% of all breast cancers in the United States, and is associated with poor prognosis (Yin et al., 2020).

Several factors can increase the risk of developing breast cancer, and this includes modifiable risk factors such as obesity (Lauby-Secretan et al., 2016). The role of obesity in breast cancer has received significant attention in recent years, in part owing to the increased prevalence of obesity in general, with half of adults globally and 70% of adults in the United States being considered overweight or obese (Gjermeni et al., 2021). Obesity significantly increases the risk of postmenopausal breast cancer, with each 5-unit increase in BMI linked to a 12% increase in risk (Renehan et al., 2008). In addition, obesity increases the risk of large, high-grade tumors, metastasis, and recurrence regardless of menopausal status (Calle and Kaaks, 2004; Carmichael, 2006; Ewertz et al., 2011; Kuziel et al., 2023; Mazzarella et al., 2013). Obesity is a risk factor for both estrogen receptor-positive and TNBC, suggesting that a multitude of mechanisms underlie the obesity-breast cancer link (Berger and Iyengar, 2021).

Obesity is characterized by an expansion of white adipose tissue, the major tumor microenvironment of breast cancer. Thus, there is significant interest in elucidating how the cellular constituents of adipose tissue may accelerate breast cancer cell growth. White adipose tissue is comprised of energy-storing endocrine cells called adipocytes, adipose stem and progenitor cells (ASPC), endothelial cells, and immune cells (Emont et al., 2022). The role of adipocytes in promoting breast cancer has received considerable attention in recent years. Cancer-associated adipocytes were shown to release adipokines, chemokines, growth factors, and extracellular matrix proteins that contribute towards tumor development and metastasis (Wu et al., 2019; Zhao et al., 2020). Several studies have highlighted the importance of the ratio of two adipokines in particular, leptin and adiponectin, to the progression of postmenopausal breast cancer as well as TNBC (Chen et al., 2006; Grossmann et al., 2008; Sultana et al., 2017). Adiponectin, whose expression is decreased with obesity, inhibits breast cancer growth, though this action depends on estrogen receptor status (Musgrove and Sutherland, 2009; Naimo et al., 2020). In contrast, leptin, whose expression is increased with obesity, promotes breast cancer growth (Strong et al., 2015; Wei et al., 2016). Similarly, resistin, whose secretion is linked to obesity and diabetes, can promote breast cancer (Gao et al., 2020). Finally, Maguire et al showed that obese adipocytes synthesize and secrete increased amounts of creatine, which promotes the growth of the TNBC cell line E0771 in obese mice (Maguire et al., 2021).

The other cellular components of adipose tissue may similarly promote breast cancer growth, though the underlying mechanisms have not been described in much detail. Breast cancer promotes increased circulation and recruitment of ASPCs to the tumor microenvironment, particularly in obese patients (Zhang et al., 2009). Notably, orthotopic xenografts of breast cancer cells in combination with ASPCs show increased tumor growth and metastasis, further enhanced if ASPCs were isolated from obese individuals (Chandler et al., 2012; Hillers et al., 2018). Depletion of ASPCs in the microenvironment inhibits tumorigenesis (Daquinag et al., 2016). How ASPCs promote breast cancer growth remains unclear. Possible mechanisms include differentiating into additional cancer-associated adipocytes, secreting angiogenic or inflammatory factors, or transdifferentiating into fibroblast-like cells to increase tumor microenvironment stiffness (Kuziel et al., 2023; Zhang et al., 2012). Not all of these mechanisms are mutually exclusive and the relative contributions of these mechanisms to breast cancer growth may depend on clinical subtype and tumor grade. We do not yet know how breast cancer may regulate the cell fate and function of ASPCs.

Fibrosis describes the stiffening of tissue due to the deposition of extracellular matrix (ECM) components such as collagen (Wells, 2022). Studies have shown that breast cancer cells are more proliferative and invasive when grown *ex vivo* in stiff ECM and *in vivo* when injected into fibrotic tissue (Barcus et al., 2017; Levental et al., 2009; Provenzano et al., 2008; Shea et al., 2018). Accordingly, increased fibrosis in the tumor microenvironment is linked to increased breast cancer aggressiveness (Kuziel et al., 2023). Cancer-associated fibroblasts (CAFs) are thought to be the main culprit of fibrosis in the tumor microenvironment. However, recent single-cell sequencing has shown that there are multiple subtypes of CAFs, potentially reflecting different origins (Bartoschek et al., 2018). This includes mature adipocytes, which were shown capable of becoming more fibroblast-like in the presence of breast cancer cells *ex vivo* (Bochet et al., 2013). ASPCs are another source of CAFs, and recent single-cell transcriptomic analyses have identified ASPC subpopulations with fibroblast gene expression signatures (Hepler et al., 2018; Sarvari et al., 2021). Notably, ASPCs isolated from obese adipose tissue or from adipose tissue adjacent to breast cancer show increased fibroblast-like gene expression, likely contributing to the increased amounts of adipose tissue fibrosis observed in both settings (Cozzo et al., 2017; Kidd et al., 2012; Miran et al., 2020; Sarvari et al., 2021; Yang et al., 2023). The mechanisms underlying this pathogenic remodeling and how breast cancer cells may instruct the fate of ASPCs in the surrounding adipose tissue toward fibroblasts remain unclear.

In this study, we screened the secretomes of ten human breast cancer cell lines, encompassing all clinical subtypes, for the ability to modulate adipogenic differentiation of ASPCs. The secretomes of all TNBC cell lines tested potently inhibited adipocyte differentiation. Using mass spectrometry, we identified that a subset of TNBC cells secrete Zinc Alpha-2-Glycoprotein, ZAG/AZGP1, an adipokine previously linked to adipocyte lipolysis and cancer cachexia (Bing and Trayhurn, 2009). We show that ZAG treatment of ASPCs promotes the expression of fibrotic genes. Accordingly, loss of ZAG dramatically reduces fibrosis in the tumor microenvironment and impedes tumor growth.

## Results

### Breast cancer cells secrete factors that modulate adipogenesis

Previous studies have shown that breast cancer cells can transform the white adipose tissue tumor microenvironment to become more tumor supportive (Nguyen et al., 2021; Tang et al., 2023). To identify factors secreted by breast cancer cells to regulate ASPC fate, we obtained 10 distinct human breast cancer cell lines, encompassing all clinical subtypes (Figure S1A). Once breast cancer cell lines were confluent, the growth media was changed to defined Dulbecco’s Modified Eagle Medium (DMEM) only media. Conditioned media containing breast cancer cell secretome was collected 72 hours later. To study any potential effect of the secretome on adipogenesis, we used a well-characterized, murine cell line model of adipogenesis, 3T3-L1 cells (Green and Meuth, 1974). These cells differentiate uniformly in response to an adipogenic differentiation cocktail containing insulin, dexamethasone, and the cAMP-elevating agent 3-isobutyl-1-methylxanthine (IBMX).

3T3-L1 cells were differentiated using breast cancer cell conditioned media mixed in a one-to-one ratio with fresh DMEM and supplemented with serum and adipogenic differentiation cocktail (Figure 1A). 3T3-L1 adipogenesis was monitored by quantifying intracellular lipid accumulation using a lipophilic, green fluorescent dye BODIPY 493/503 and the IncuCyte live-cell imaging system. This allows for both quantification of intracellular lipid content at the end-point of the adipogenesis protocol and kinetic analysis of intracellular lipid accumulation during adipogenesis. Strikingly, the secretome of most breast cancer cells significantly modulated 3T3-L1 adipogenesis (Figure 1B, S1B, C). In particular, the secretomes of all TNBC cells tested severely inhibited 3T3-L1 adipogenesis (Figure 1B, S1B, C). Only the secretome of the luminal A breast cancer cell line MCF7 promotes 3T3-L1 adipogenesis. The secretomes of luminal B breast cancer cell lines and HER2-positive breast cancer cell lines had more modest or no effects on 3T3-L1 adipogenesis.

**Figure 1:**
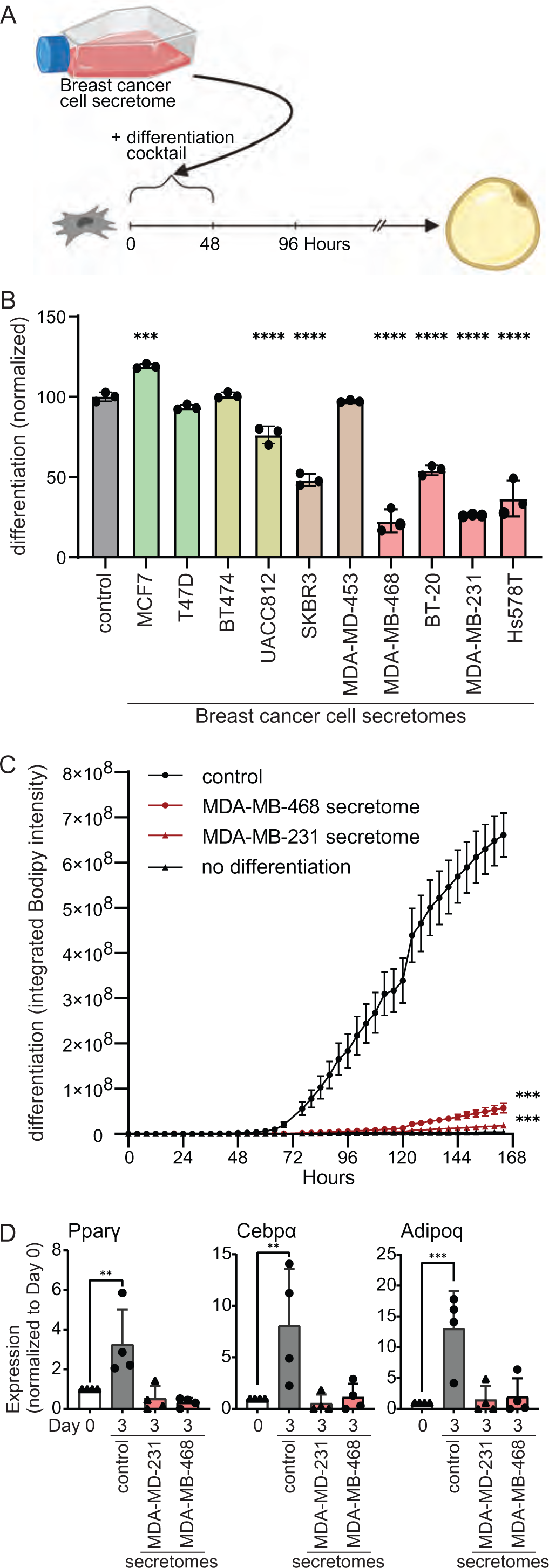
Secretome of human breast cancer cells modulates adipogenesis. (A) Schematic of experimental workflow. Breast cancer cell secretomes were added only during the first two days of 3T3-L1 adipogenesis. (B) Effect of human breast cancer cell line secretomes on 3T3-L1 adipogenesis. (C-D) The secretome of TNBC cell lines MDA-MB-468 and MDA-MB-231 inhibits the differentiation of primary murine ASPCs as assessed by (C) lipid accumulation during adipogenesis and (D) expression of adipogenic genes *Ppar*γ, *Cebp*α, and *Adipoq*. (B-D) All data are mean ± SD. Data points show independent biological replicates. p-values calculated using (B, D) one-way ANOVA followed by Dunnett’s multiple comparison test or (C) two-way ANOVA followed by Šidák’s multiple comparison test. (**<0.01, ***<0.001 and ****<0.0001). See also Figure S1.

Given the dramatic and uniform effect of TNBC secretomes on 3T3-L1 adipogenesis, we next assessed its effect on primary ASPCs freshly isolated from the inguinal white adipose tissue of C57Bl/6 mice. Remarkably, the presence of TNBC secretome completely abrogated adipogenesis as assessed by intracellular lipid content (Figure 1C, S1D). Similarly, the presence of TNBC secretome inhibited the expression and activation of known adipogenic transcription factors and regulators as assessed by quantitative real-time PCR (Figure 1D) or using a PPARγ reporter cell line (Hilgendorf et al., 2019) (Figure S1E). Thus, all TNBC cell lines tested secrete one or more paracrine factors that potently inhibit the differentiation of ASPCs.

### Zinc Alpha-2-Glycoprotein (ZAG) is necessary and sufficient for TNBC-mediated inhibition of adipogenesis

To identify the anti-adipogenic factor secreted by TNBC cells, we first sought to determine the class of macromolecule. Specifically, we heat inactivated the secretome of two TNBC cell lines, MDA-MB-231 and MDA-MB-468 cells, to denature heat-labile proteins. Heat inactivation completely rescued 3T3-L1 adipogenesis (Figure 2A, S2A). Thus, the secretome of these two TNBC cell lines contains one or more heat-labile proteins required for the inhibition of adipogenesis. Next, we sought to enrich for a fraction containing this anti-adipogenic protein(s) using centrifugal filter columns. TNBC secretomes were subjected to spin columns with a molecular weight cut-off of 100,000 Daltons. 3T3-L1 cells were then differentiated in the presence of either the filtrate or supernatant. The ability of the TNBC secretome to inhibit 3T3-L1 adipogenesis was only retained in the supernatant (Figure 2B, S2B). Of note, the conditioned media containing TNBC secretome was collected in defined DMEM only as described above (i.e., no serum), which does not contain any proteins greater than 100,000 Daltons in size. Thus, our initial biochemical characterization of the TNBC secretome likely yielded a fraction relatively depleted of proteins but highly enriched for the unknown anti-adipogenic protein(s) of interest.

**Figure 2:**
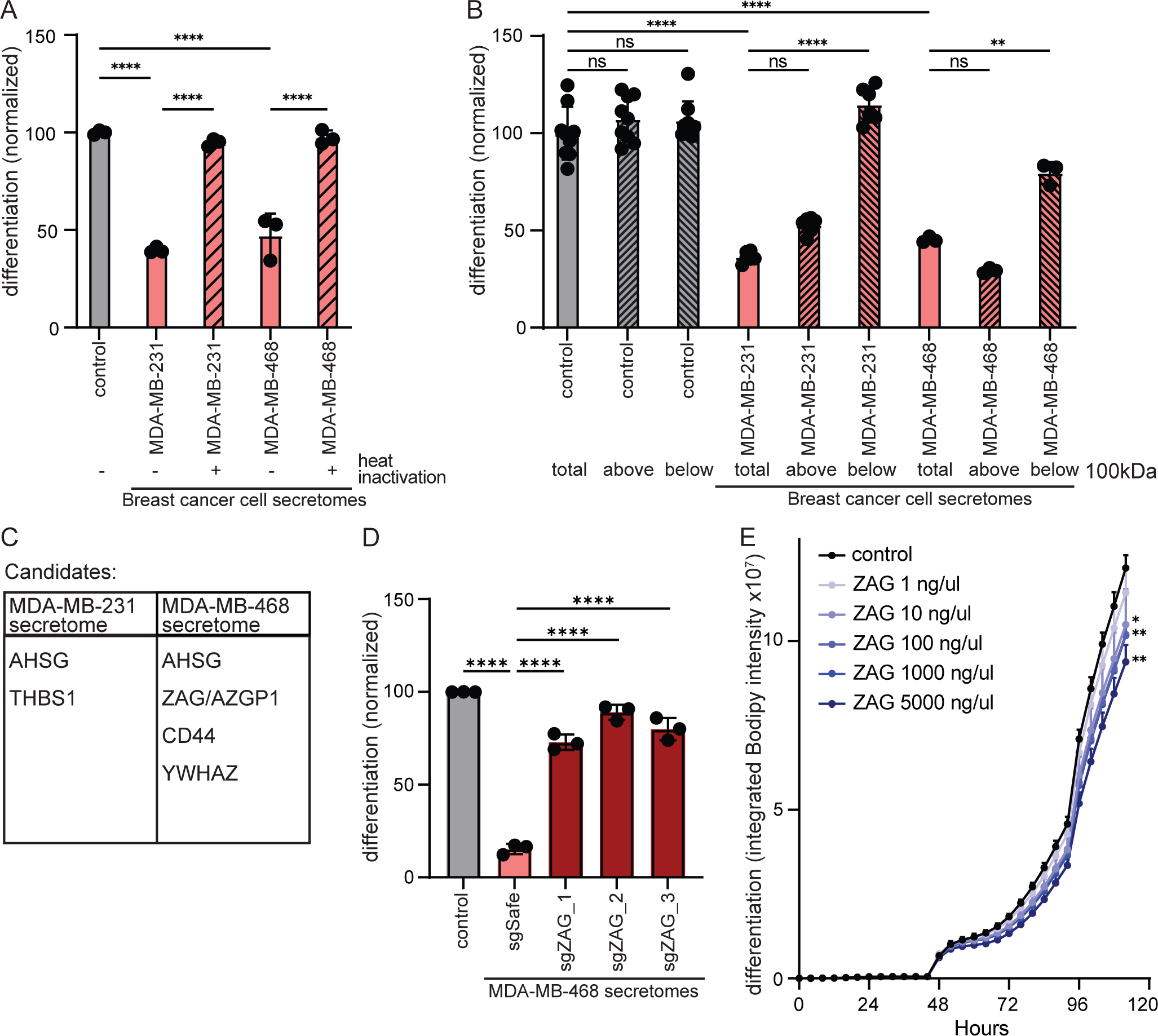
The glycoprotein ZAG inhibits adipogenesis. (A) Inhibition of 3T3-L1 adipogenesis is rescued with heat inactivation of the secretome of MDA-MB-468 and MDA-MB-231 cells. (B) The anti-adipogenic factor of the MDA-MB-468 and MDA-MB-231 secretomes (total) is retained in the supernatant (top) fraction following centrifugation using a spin column with a molecular weight cut-off of 100,000 Daltons. (C) Candidate anti-adipogenic factors identified in the secretome of MDA-MB-468 and MDA-MB-231 cells. (D) The secretome of MDA-MB-468 cells depleted of ZAG does not inhibit 3T3-L1 adipogenesis. (E) Supplementing 3T3-L1 cells with recombinant ZAG during the first two days of adipogenesis is sufficient to inhibit adipogenesis. (A-E) All data are mean ± SD. Data points show independent biological replicates. p-values calculated using one-way ANOVA followed by (A-B) Tukey’s multiple comparison test or (D) Dunnett’s multiple comparison test, or (E) two-way ANOVA followed by Šidák’s multiple comparison test. (ns is non-significant; *<0.05, **<0.01, and ****<0.0001). See also Figure S2.

We next performed protein mass spectrometry on the spin column supernatants of three independent secretomes collected from two TNBC cell lines, MDA-MB-231 and MDA-MB-468 cells (Table S1). After excluding abundant intracellular protein contaminants, this analysis yielded five candidate anti-adipogenic proteins, only one of which was found in the secretomes of both TNBC cell lines (Figure 2C). To identify which of these candidate anti-adipogenic proteins is required in the TNBC secretome to inhibit 3T3-L1 adipogenesis, we next individually knocked out each candidate in the corresponding TNBC cell line using Crispr/Cas9. Specifically, we generated three MDA-MB-468 cell lines each depleted for ASHG, ZAG (also known as AZGP1), CD44, or YWHAZ. Remarkably, the secretome of all three MDA-MB-468 cell lines depleted for ZAG rescued 3T3-L1 adipogenesis (Figure 2D, S2C, D). Thus, the presence of ZAG is required for the MDA-MB-468 secretome to inhibit 3T3-L1 adipogenesis. Loss of AHSG, CD44, or YWHAZ did not rescue adipogenesis, nor was recombinant TSP-1 sufficient to inhibit 3T3-L1 adipogenesis (Figure S2E, F). In contrast, the presence of recombinant ZAG during the first two days of 3T3-L1 adipogenesis is sufficient to inhibit adipogenesis in a dose-dependent manner, albeit not as efficiently as the MDA-MB-468 secretome (Figure 2E, S2G). Together, these data show that ZAG is both necessary and sufficient for MDA-MB-468 cells to inhibit adipogenesis. Of note, ZAG was not identified by proteomics or immunoprecipitation in the secretome of MDA-MB-231 cells (Table S1, Figure S3A). Since the MDA-MB-231 secretome also potently inhibits adipogenesis, this suggests either that MDA-MB-231 cells secrete another large and heat-labile protein capable of inhibiting adipogenesis, or that any ZAG secreted by MDA-MB-231 is below our detection limits due to technical limitation.

### ZAG is expressed by a subset of TNBC cells and linked to poor prognosis

Zinc Alpha-2-Glycoprotein (ZAG) is primarily secreted by mature adipocytes and promotes adipocyte lipolysis to enable energy homeostasis (Banaszak et al., 2021). Plasma levels in healthy adults are 18-30mg/dl, and this is decreased in obese patients and mice (Drozdz et al., 2021; Wang et al., 2020). Of note, the level of ZAG present in the MDA-MB-468 secretome capable of inhibiting adipogenesis is therefore well within the physiologically relevant range. (Figure S3B). Interestingly, ZAG expression is increased in the adipose tissue of patients with cancer cachexia, and ZAG is identical to the lipid-mobilizing factor (LMF) purified from urine of patients with cancer cachexia (Todorov et al., 1998). Moreover, ZAG has also been shown to be expressed by several types of malignant tumors, including breast, prostate, and lung (Albertus et al., 2008; Bondar et al., 2007; Diez-Itza et al., 1993; Hale et al., 2001).

ZAG secreted by adipocytes or cancer cells is known to be highly glycosylated (Liu et al., 2020; Yan et al., 2024). Accordingly, we observed the presence of high molecular weight species in the secretome of MDA-MB-468 parental cells and control cells expressing an sgRNA targeting a safe locus (Figure 3A). These high molecular weight species are only present in the secretome, not the cell lysate, and are greatly depleted in a pool of MDA-MB-468 cells depleted of ZAG, confirming specificity of the antibody.

**Figure 3:**
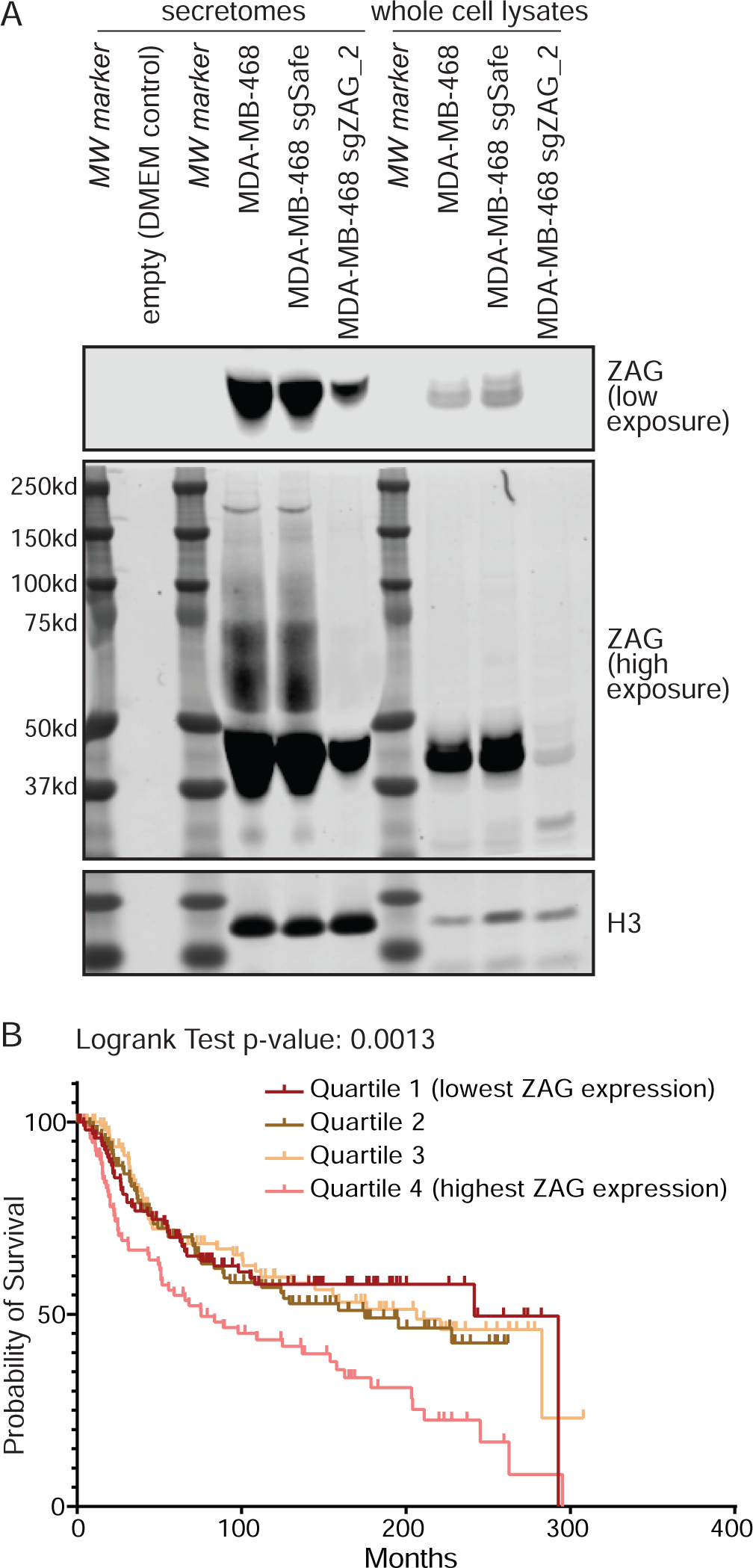
ZAG is secreted by TNBC cells and linked to poor prognosis. (A) MDA-MB-468-secreted ZAG is highly modified (lanes 4, 5). These high molecular weight species were not observed in the secretome of ZAG depleted cells (lane 6), DMEM-only media control (lane 2), or in whole cell MDA-MB-468 lysates (lanes 8, 9). Histone 3 used as lysate loading control and also found in secretome, likely due to sparse cell death. (B) TNBC outcome of combined Metabric and TCGA data sets stratified by ZAG expression. See also Figure S3.

Next, we analyzed two large breast cancer patient gene expression data sets (TCGA (Cancer Genome Atlas, 2012) and Metabric (Curtis et al., 2012; Pereira et al., 2016; Rueda et al., 2019)) using cBioPortal (Cerami et al., 2012; de Bruijn et al., 2023; Gao et al., 2013). Consistent with our inability to detect ZAG in the secretome of MDA-MB-231 cells, this analysis shows that ZAG expression in TNBC cells is highly variable (Figure S3C). Remarkably, stratifying clinical outcome of all TNBC by ZAG expression shows that patients with TNBC expressing ZAG at high levels have significantly worse prognosis (Figure 3B, S3D). This suggests that the ZAG secreted by TNBC cells and capable of inhibiting adipogenesis promotes tumorigenesis. Interestingly, the mean expression of ZAG is lower in TNBC than in estrogen receptor (ER)- positive breast cancer (Figure S3C). However, stratifying clinical outcome of ER-positive breast cancer by ZAG expression shows no correlation with poor prognosis (Figure S3E, F). Thus, our study identified a potential TNBC-specific function of ZAG in promoting tumorigenesis.

### ZAG promotes TNBC tumor growth and induces the expression of fibrotic genes in ASPCs

To explore the role of ZAG in MDA-MB-468 tumorigenesis, we next injected control or ZAG depleted cells into the 4^th^ mammary fat pad of female NRG mice. ZAG depletion potently inhibits orthotopic xenograft growth (Figure 4A, S4A). Consistent with decreased tumor burden at the experimental end point (Figure 4B, S4B), loss of ZAG resulted in decreased proliferation of MDA-MB-468 cells *in vivo*, though this difference did not reach significance (Figure 4C). Importantly, these effects were not cell intrinsic, since loss of ZAG does not affect MDA-MB-468 proliferation *ex vivo* (Figure S4C).

**Figure 4:**
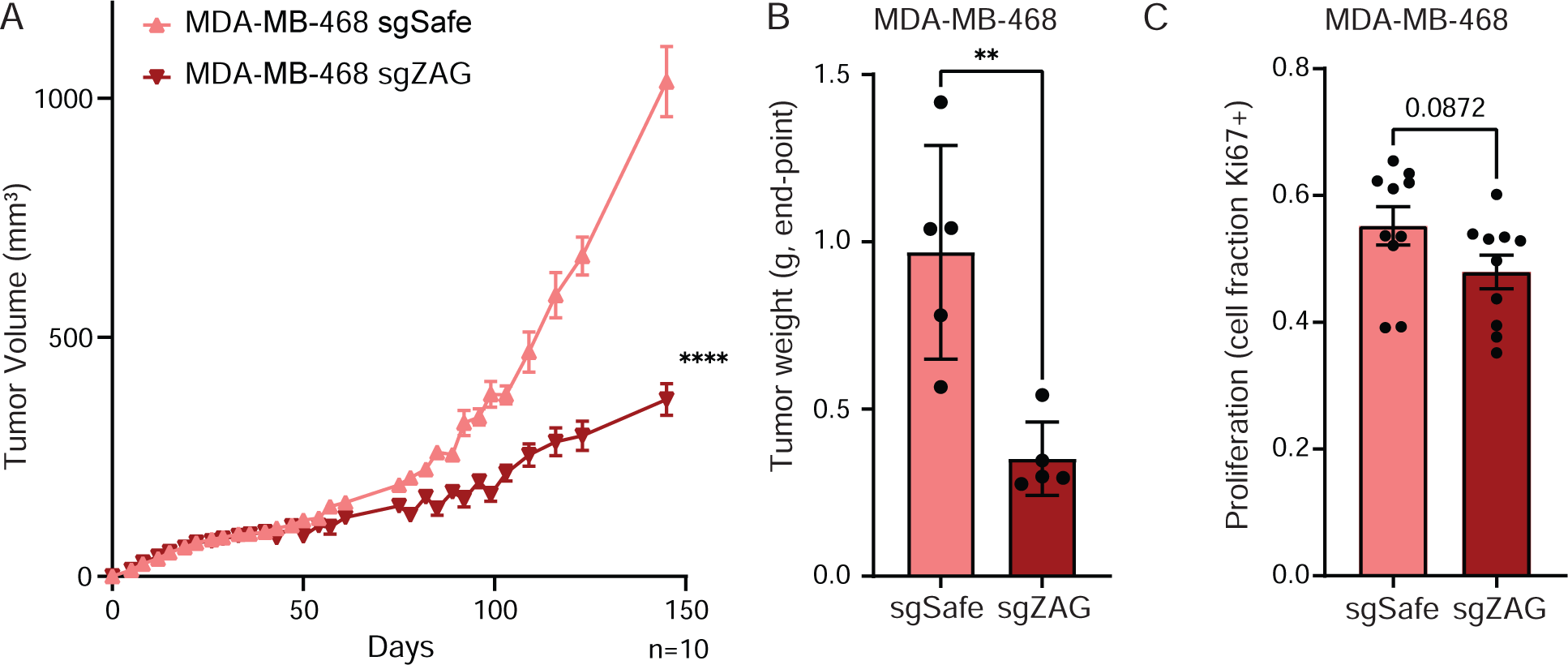
ZAG is important for MDA-MB-468 tumorigenesis. (A-B) Depletion of ZAG inhibits xenograft growth of MDA-MB-468 cells as determined by (A) tumor volume measurements over time or (B) weight of excised tumor at end-point. (C) MDA-MB-468 cells depleted for ZAG show a trend of decreased proliferation *in vivo*. (A-C) Data are mean ± SEM. Data points and n are independent mice. p-values calculated using (A) two-way ANOVA followed or (B-C) Student’s t-test. (**<0.01 and ****<0.0001). See also Figure S4.

Given the known role of ZAG in inducing lipolysis, we first quantified the size of tumor-adjacent adipocytes. Loss of ZAG did not affect the size of cancer-associated adipocytes (Figure 5A, S5A-B). Since we had initially identified ZAG as a factor that inhibits ASPC adipogenesis, we next hypothesized that it may instead promote the transdifferentiation of ASPCs into cancer-associated fibroblasts. Remarkably, staining the adipose tissue tumor microenvironment for markers of fibrosis uncovered dramatic differences. Specifically, ZAG depletion in MDA-MB-468 cells caused a significant reduction in both Picro Sirius Red staining and Alpha-Smooth Muscle Actin (αSMA) staining in the surrounding adipose tissue (Figure 5B-D, S5C). We therefore assayed for the induced expression of fibrotic markers in primary murine ASPCs differentiated in the presence of MDA-MB-468 secretome. Consistent with the increased fibrosis observed in the adipose tissue tumor microenvironment, we show that exposing primary ASPCs to the secretome of MDA-MB-468 cells promotes the expression of canonical fibrosis genes (Figure 5E). Thus, we identified a glycoprotein, ZAG, that is secreted by a subset of TNBC cells and may promote tumorigenesis by inhibiting adipogenesis and instead promoting the transdifferentiation of ASPCs into cancer-associated fibroblasts.

**Figure 5:**
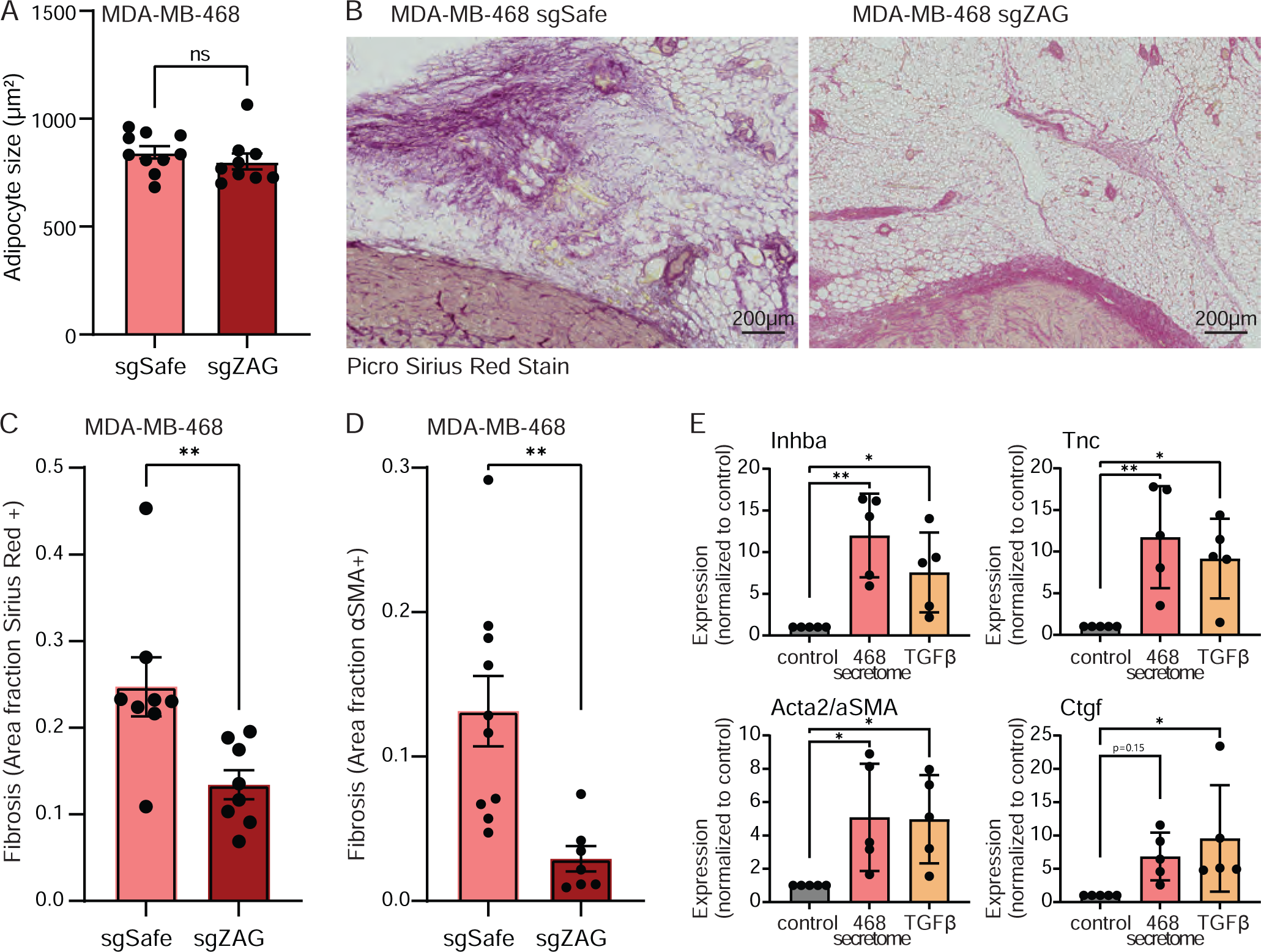
TNBC-secreted ZAG promotes fibrosis in the surrounding adipose tissue. (A) The average size of cancer-associated adipocytes is not affected by ZAG depletion in MDA-MB-468 xenografts. (B-D) The adipose tissue microenvironment of MDA-MB-468 xenografts depleted for ZAG are less fibrotic. (B) Representative image and (C) quantification of Picro Sirius Red staining. (D) Quantification of αSMA staining. (D) The secretome of MDA-MB-468 cells promotes the expression of fibrotic genes *Inhba*, *Tnc*, *Acta* (α*Sma*), and *Ctgf* in differentiating primary murine ASPCs (day 3 of adipogenic differentiation). TGFβ treatment of ASPCs for three days shows similar induced expression of fibrotic genes. (A-D) Data are mean ± SEM. Data points are quantification of the staining of sections from independent mice. p-values calculated using Student’s t-test. (E) Data are mean ± SD. Data points show independent biological replicates from separate ASPC isolations. p-values calculated using one-way ANOVA followed by Dunnett’s multiple comparison test. (ns is non-significant; *<0.05 and **<0.01). See also Figure S5.

## Discussion

The adipose tissue tumor microenvironment is critical for breast cancer growth, and breast cancer cells are known to transform the surrounding adipose tissue into a more tumor-supportive microenvironment (Nguyen et al., 2021; Tang et al., 2023; Trivanovic et al., 2020). Here, we screened the secretomes of 10 different human breast cancer cell lines, encompassing all clinical subtypes, for the ability to regulate ASPC fate. We identified the cachexia-associated glycoprotein, ZAG, which promotes TNBC growth and fibrosis in the adipose tissue tumor microenvironment. High ZAG expression in TNBC is linked to poor prognosis. Together, these data suggest that targeting ZAG in TNBC may have therapeutic potential.

In healthy adults, ZAG is primarily secreted by mature adipocytes to promote lipid mobilization in an autocrine or paracrine manner (Bao et al., 2005; Bing et al., 2010). Previous studies have shown that ZAG is also expressed by malignant tumor cells, including breast cancer cells (Diez-Itza et al., 1993; Dubois et al., 2010). Xenograft studies using ZAG-overexpressing cells show shrunken, beige fat-like adipocytes (Elattar et al., 2018). Overexpression of ZAG in 3T3-L1 adipocytes inhibits the expression of lipogenic genes, and stimulates adipocyte lipolysis and beiging through increased expression of lipases, including adipose triglyceride lipase (ATGL) and hormone-sensitive lipase (HSL), and beige fat-specific markers (Xiao et al., 2018; Zhu et al., 2013). Here, we identified a novel function of ZAG in regulating ASPC fate in adipose tissue. This discovery is distinct and separate from lipogenesis/lipolysis in mature adipocytes, since we observe a decrease in the expression of canonical adipogenic transcription factors when differentiation of ASPCs is initiated in the presence of ZAG. Further, we do not observe any changes in adipocyte size in the tumor microenvironment in xenograft studies using ZAG-depleted breast cancer cells. Together, this suggests that TNBC cells may use ZAG to remodel the adjacent adipose tissue to both increase stiffness by targeting ASPCs and/or increase lipid fuels by targeting mature adipocytes. Follow-up investigation will establish how TNBC-secreted ZAG promotes the trans-differentiation of ASPCs into fibroblast-like cells.

We show here that high ZAG expression by TNBC is associated with poor prognosis. However, our comprehensive analysis of breast cancer patient gene expression data did not reveal any unidirectional association between ZAG expression and prognosis when all clinical subtypes of breast cancer were combined. Elevated ZAG expression may even be associated with better prognosis in ER-positive breast cancer. Interestingly, while ZAG expression is significantly elevated in breast cancer tissue compared to normal breast tissue from healthy women, previous studies have linked ZAG expression in breast cancer with both worse and better prognoses (Diez-Itza et al., 1993; Dubois et al., 2010; Parris et al., 2014; Qin et al., 2024). The clinical subtype of breast cancer was not considered in most of these studies, potentially accounting for these disparate observations. Mechanistically, depletion of ZAG in two human breast cancer cell lines, MCF7 and MDA-MB-231, was shown to inhibit proliferation, migration, and invasion *ex vivo*, and tumor growth *in vivo* (Qin et al., 2024). Recombinant ZAG was also shown to promote the proliferation of MCF7 and MDA-MB-231 cells (Delort et al., 2013). We note that we could not detect ZAG in MDA-MB-231 cells (Figure S3A), consistent with no noted expression in publicly available gene expression data (Jin et al., 2023) [proteinatlas.org], and did not observe a cell-intrinsic proliferation defect in MDA-MB-468 cells depleted of ZAG (Figure S4C) in our study. Instead, we show here that recombinant ZAG and MDA-MB-468 secreted ZAG inhibit adipogenesis and promote adipose tissue fibrosis and tumor growth.

Beyond breast cancer, ZAG expression is significantly elevated in colorectal cancer compared to normal tissue, and high ZAG expression or elevated serum ZAG levels are linked with poor prognosis in colorectal cancer (Fang et al., 2022; Ji et al., 2013; Liu et al., 2022). ZAG expression is further elevated in colorectal cancer with liver metastasis compared to colorectal cancer without distant metastasis, and ZAG was shown to promote epithelial-to-mesenchymal trans-differentiation and modulate the expression adhesion proteins (Ji et al., 2019). In contrast, reduced ZAG expression is linked to poor prognosis of prostate cancer, gastric cancer, and hepatocellular carcinoma (Brooks et al., 2016; Huang et al., 2013; Huang et al., 2012). ZAG has also been shown to suppress the growth of oral and pancreatic cancer by downregulating cyclin-dependent kinase 1 gene and inducing mesenchymal-to-epithelial transdifferentiation, respectively (He et al., 2001; Kong et al., 2010). Beyond cancer, ZAG was shown to inhibit fibrosis in the kidney and heart (Sorensen-Zender et al., 2015). Together, these findings reveal that ZAG exhibits distinct and often opposing functions depending on the cellular context of its expression.

We propose that ZAG secreted by a subset of TNBC is functionally distinct compared to ZAG secreted by ER-positive breast cancer. Specifically, we show here that TNBC-secreted ZAG promotes the trans-differentiation of ASPCs into cancer-associated fibroblasts. We do not yet know how ZAG secreted by other breast cancer subtypes regulates ASPC fate. How could TNBC-secreted ZAG have a distinct function? One possibility is that ZAG, a highly glycosylated protein, is differentially glycosylated by TNBC cells, and that only the TNBC-specific glycosylation pattern enables ZAG to regulate ASPC fate. Of note, ZAG is abundant in human saliva and a potential biomarker for lung cancer (Xiao et al., 2012); LC-MS/MS analysis of salivary ZAG identified 22 glycan structures, five of which were unique to samples collected from lung cancer patients (Liu et al., 2020). We do not yet know the glycan structure of TNBC-secreted ZAG. Alternatively, other paracrine factors secreted by TNBC cells may be required for ZAG to promote ASPC trans-differentiation and fibrosis. We note that either model may explain why the recombinant ZAG used in this study was not as efficient at inhibiting adipogenesis as TNBC-secreted ZAG. Future investigation will establish if and how TNBC-secreted ZAG differs from ZAG secreted by other sources in terms of posttranslational modifications or available interacting proteins. Since ZAG expression is linked with poor or better prognosis across several types of cancer (Brooks et al., 2016; Huang et al., 2013; Huang et al., 2012; Ji et al., 2019), there is also a need to broadly interrogate the function of ZAG sourced from a multitude of types of cancers.

Even within TNBC, we found significant heterogeneity in ZAG expression. Since TNBC is defined as a lack of markers, this likely reflects the inherent histological and molecular heterogeneity of cancers classified as TNBC. Nonetheless, all four TNBC secretomes tested inhibited adipogenesis. Our follow-up investigation on two of these cell lines, MDA-MB-468 and MDA-MB-231, revealed that ZAG is only expressed by MDA-MB-468 cells. We do not yet know how MDA-MB-231 cells inhibit adipogenesis. MDA-MB-231 cells may secrete a different heat-labile, large factor, or we were merely unable to detect ZAG in the MDA-MB-231 secretome due to technical limitations, such as low expression, differential modifications, or a limited understanding of how ZAG expression is regulated. Of note, one recent study showed that ZAG expression is directly regulated by androgen signaling, and that ZAG is specifically secreted by androgen-responsive TNBC, patient-derived xenograft models (Hanamura et al., 2021). Thus, specific subtypes within TNBC may modulate the tumor microenvironment via secreted ZAG.

Finally, obesity is a known risk factor for breast cancer, including TNBC (Sun et al., 2017). Understanding the underlying mechanisms is complicated by the multifactorial effects of obesity, such as alterations to the cellular composition and morphology of adipose tissue, hyperglycemia, hyperinsulinemia, and hyperlipidemia. Obesity is linked to increased fibrosis in adipose tissue, which is thought to promote breast cancer cell growth. We do not yet know how obesity affects ZAG expression by TNBC, ZAG-dependent remodeling of the adipose tissue tumor microenvironment, and ZAG expression-linked patient prognosis.

## Acknowledgments

This work was supported by grants from 5 For The Fight and Huntsman Cancer Institute (KIH), the V Foundation for Cancer Research (KIH), and the Pew Charitable Trusts (KIH). Proteomics mass spectrometry analysis was performed at the Mass Spectrometry and Proteomics Core Facility at the University of Utah. Mass spectrometry equipment was obtained through a Shared Instrumentation Grant 1 S10 OD018210 01A1. Research reported in this publication utilized the Preclinical Research Shared Resource and the Cancer Bioinformatics Shared Resource at Huntsman Cancer Institute at the University of Utah and was supported by the National Cancer Institute of the National Institutes of Health under Award Number P30CA042014. The content is solely the responsibility of the authors and does not necessarily represent the official views of the NIH. We acknowledge the Cell Imaging Core at the University of Utah for use of equipment (Zeiss Axio Scan.Z1 Slide Scanner) and the Flow Cytometry Core at the University of Utah for use of equipment (BD FACSAria). Graphical abstract created with BioRender.com

## Author Contributions

Conceptualization, SV and KIH; Methodology, SV, SDG, AA, DHL, JEC, and KIH; Software, SV and SMO; Investigation, SV, SDG, MCC, MA, SMO, GW; Resources, DHL, JEC, and KIH; Writing – Original Draft, SV and KIH; Writing – Review & Editing, SV, SDG, and KIH; Supervision, DAN, DHL, JEC, and KIH; Funding Acquisition, KIH.

## Supplemental Information titles and legends

**Figure S1: related to Figure 1. Breast cancer line secretomes modulate adipogenesis.**

(A) Table of human breast cancer cell lines used, categorized according to clinical subtype noted by the American Type Culture Collection. (B) Time course of lipid accumulation as assessed by total integrated green fluorescent BODIPY intensity during 3T3-L1 adipogenesis. See Figure 1B for endpoint analysis of adipogenesis. The secretomes of human breast cancer cell lines modulates adipogenesis. (C) Images showing total lipid accumulation of 3T3-L1 cells differentiated in the presence of human breast cancer cell line secretome using Oil Red O staining at experimental endpoint. (D) End-point analysis of total lipid accumulation as a measure of adipogenesis. See Figure 1C for time course analysis of lipid accumulation during adipogenesis. The secretomes of TNBC cell lines MDA-MB-468 and MDA-MB-231 inhibit the differentiation of primary murine ASPCs as assessed by total integrated green fluorescent BODIPY intensity at the end-point of 3T3-L1 adipogenesis and normalized to DMEM control (no secretome). (E) A PPARγ-T2A-GFP knockin reporter shows decreased expression of PPARγ in 3T3-L1 cells differentiated in the presence of the secretome of MDA-MB-468 and MDA-MB-231 cells. (B) Data are represented as mean ± SD. (D) Data are represented as mean ± SD. p-values calculated using one-way ANOVA followed by Dunnett’s multiple comparison test (****<0.0001). (E) Data are represented as mean ± SD. p-values calculated using two-way ANOVA followed by Šidák’s multiple comparison test (***<0.001, ****<0.0001)

**Figure S2: related to Figure 2. Identification of ZAG in the MDA-MB-468 secretome as an anti-adipogenic factor.**

(A) Time course of lipid accumulation. See Figure 2A for end-point adipogenesis analysis. Heat-inactivated MDA-MB-468 or MDA-MB-231 secretomes do not inhibit 3T3-L1 adipogenesis. (B)Time course analysis of lipid accumulation. See Figure 2B for end-point adipogenesis analysis. 3T3-L1 cells were differentiated in the presence of unfractionated MDA-MB-468 or MDA-MB-231 secretomes (CM), the supernatant following centrifugal filtration (above 100kd), or the filtrate (below 100kd). The top fractions of fractionated MDA-MB-468 or MDA-MB-231 secretomes inhibit 3T3-L1 adipogenesis. Of note, spin columns do not accurately fractionate all proteins according to indicated molecular weight due to differential folding and/or posttranslational modifications, and cut-off values should not be regarded as definitive values of molecular weights for unknown proteins. (C) Time course of lipid accumulation. See Figure 2D for end-point adipogenesis analysis. ZAG is required for the MDA-MB-468 secretome to inhibit 3T3-L1 adipogenesis. (D) TIDE analysis showing genomic alterations in the three engineered MDA-MB-468 sgZAG cell lines. Each cell line was infected with the one sgRNA (see Table S2) and cell lines are pools of recombination (i.e., not clonal cell lines). (E) Time course of lipid accumulation of 3T3-L1 cells supplemented with TSP-1 (THBS1) during first 2 days of adipogenesis. TSP-1 does not modulate 3T3-L1 adipogenesis. (F) The secretomes of MDA-MB-468 cells lacking YWHAZ, AHSG, or CD44 retain the ability to inhibit 3T3-L1 adipogenesis, and the presence of these candidate proteins in the secretome is not required to inhibit adipogenesis. Three distinct cell lines were generated per candidate factor. MDA-MB-468 parental cells are uninfected, MDA-MB-468 sgSafe cells are Cas9-BFP expressing cells infected with an sgRNA targeting a safe genomic locus. (G) Adipogenesis endpoint analysis of Figure 2E. Mouse myeloma NS0 cell line-derived human ZAG protein inhibits 3T3-L1 adipogenesis in a dose-dependent manner. (A-C, E) Data are represented as mean ± SD. (F-G) Data are represented as mean ± SD. p-values calculated using one-way ANOVA followed by (F) Šidák’s multiple comparison test or (G) Dunnett’s multiple comparison test. (ns is non-significant; *<0.05, **<0.01, and ****<0.0001).

**Figure S3: related to Figure 3. ZAG expression and linked prognosis by breast cancer subtypes.**

(A) Immunoprecipitation of endogenous ZAG in the secretomes of MDA-MB-468 and MDA-MB-231 cells. Anti-adipogenic effects of secretomes on 3T3-L1 cells was validated. ZAG is only found in the immunoprecipitated and input of MDA-MB-468 secretome (arrow, lanes 2 and 4), but not in the IgG pull down control (lane 5), ZAG pull-down of the MDA-MB-231 secreteme (lanes 7-10), or ZAG pull down of the DMEM only media control (lane 11). * denotes heavy chain band. (B) Quantification of ZAG abundance in MDA-MB-468 secretome by ELISA. Quantification was determined in the secretome of three distinct cell lines depleted of ZAG, MDA-MB-468 parental cells, and MDA-MB-468 sgSafe cells (control). (C) Expression of ZAG (mRNA expression z-scores relative to all samples) in all breast cancer, TNBC, or ER-positive breast cancer patients in the TCGA data set or the Metabric data set. Data analyzed using cBioPortal. Dotted lines show quartiles in violin plots. (D-F) Patient outcome stratified by ZAG expression for TCGA data set (left) or Metabric (right) separately for (D) TNBC, (E) ER-positive breast cancer, and (F) all breast cancer. Data analyzed using cBioPortal.

**Figure S4: related to Figure 4. Depletion of ZAG in MDA-MB-468 cells inhibits xenograft growth without affecting *ex vivo* proliferation.**

(A) Two independent cohorts of n=5 mice. See Figure 4A for combined data of both cohorts (n=10 mice). MDA-MB-468 cells infected with sgSafe (control) or sgZAG_2 were injected into the 4^th^ mammary fat pad of NSG female mice. One tumor per mouse. Tumor growth shown per mouse over time in cohort 1 (left) and cohort 2 (right). (B) Image of excised tumors of cohort 2. See Figure 4B for tumor weights. (C) Proliferation of MDA-MB-468 cells infected with sgSafe (control) or sgZAG_2 grown in culture (*ex vivo*). No cell-intrinsic proliferation defect was observed. (A) Lines are independent tumors/mice. P-values calculated using two-way ANOVA and followed by Šidák’s multiple comparison test for individual time point. (C) Data are represented as mean ± SD (n=3 independent experiments).

**Figure S5: related to Figure 5. Depletion of ZAG in MDA-MB-468 xenografts does not affect adipocyte size in the surrounding adipose tissue.**

(A) Histogram of cancer-associated adipocyte sizes. Adipocyte size was quantified in the adipose tissue surrounding MDA-MB-468 sgSafe or sgZAG_2 xenografts. Data points are binned quantification of adipocytes in sections from independent tumors/mice. Average adipocytes size for all 10 tumors shown in Figure 5A. (B) Representative H&E image of xenograft and surrounding adipose tissue. Adipocytes are lipid-filled cells. Adipocyte size was quantified for Figure 5A and Figure S5A. (C) Representative image showing αSMA (alpha smooth muscle actin) staining of 1 section. Quantification of sections from all 10 tumors shown in Figure 5D. Sirius Red and αSMA staining was quantified in the region of interest defined as within 2mm surrounding the tumor.

**Supplemental Table 1: Secretome proteomics**

**Supplemental Table 2: Primers used**

## Star Methods

### RESOURCE AVAILABILITY

#### Lead contact

Further information and requests for resources and reagents should be directed to and will be fulfilled by the Lead Contact, Keren Hilgendorf.

#### Materials availability

All unique/stable reagents generated in this study are available from the Lead Contact with a completed Materials Transfer Agreement.

#### Data and code availability

The published article includes all datasets generated during this study.

### EXPERIMENTAL MODEL AND SUBJECT DETAILS

#### *In vivo* animal studies

C57Bl/6J mice were purchased from Jackson Laboratory (000664). All animals were treated in accordance with the University of Utah, Institutional Animal Care and Use Committee guidelines and policies. Male mice between 6-8 weeks of age were used for isolation of murine primary preadipocytes. Female NRG mice at 5 weeks of age were used for the orthotopic xenograft experiments.

#### Cell line models

All breast cancer cell lines were purchased from ATCC and cultured in DMEM containing 10% FBS, 1% Pen/Strep, and 1% GlutaMAX until confluency was reached, at which point the growth media was replaced with DMEM only to collect conditioned media

3T3-L1 cells were cultured in DMEM medium containing 10% Bovine Calf Serum, 1% Pen/Strep, and 1% GlutaMAX and switched to DMEM containing 10% FBS, 1% Pen/Strep, and 1% GlutaMAX during adipogenesis.

Murine primary preadipocytes were isolated from inguinal white adipose tissue from C57Bl/6J male mice Primary preadipocytes were maintained and differentiated in DMEM medium containing 10% FBS, 1% Pen/Strep, and 1% GlutaMAX.

### METHOD DETAILS

#### Plasmids

pMCB306 (lentiviral vector, loxP-mU6-sgRNAs-puro resistance-EGFP-loxP) and p293 Cas9-BFP were gifts from Prof. Michael Bassik (Stanford University). pCMV-VSV-G and pCMV-dR8.2 dvpr were gifts from Bob Weinberg (Addgene plasmid #8454; http://n2t.net/addgene:8454; RRID:Addgene_8454 and #8455;; http://n2t.net/addgene:8455; RRID:Addgene_8455) (Stewart et al., 2003). Lentiviral vectors containing sgRNA were generated by ligating annealed sgRNA oligonucleotides into pMCB306 vector digested with BstXI and BlpI restriction enzymes.

#### Cell Line generation

Lentiviral vectors carrying the gene of interest were co-transfected with pCMV-VSV-G and pCMV-dR8.2 dvpr into 293T cells using Fugene6 (Promega). Media was replaced after 24h and virus was harvested 48 and 72h post-transfection. Virus was filtered with a 0.45μm OVDF filter (Millipore) or freeze-thawed and subjected to centrifugation at 3000rpm for 10min. Cell lines were infected with virus in 10μg/ml polybrene (Millipore). Media was replaced after 24h and infected cells were isolated via FACS sorting after 48-72h post-infection.

MDA-MB-468 cells expressing Cas9-BFP were generated by infection of virus harvested from 293T cells transfected with p293 Cas9-BFP, pCMV-VSV-G and pCMV-dR8.2 dvpr. MDA-MB-468 Cas9-BFP cells were sorted for BFP positivity. To generate Crispr/Cas9 knockout cells, MDA-MB-468 Cas9-BFP cells were infected with lentivirus containing the sgRNA of interest. Primers listed in Table S2. Knockout efficiency was determined 10 days post-infection by TIDE analysis (Brinkman et al., 2014). Cells expressing a safe-targeting sgRNA were used as control (Morgens et al., 2017). Primers listed in Table S2.

#### Breast cancer cell conditioned media

Breast cancer cells were seeded at ∼ 30 % confluency in DMEM + 10% Fetal Calf Serum + 1% Pen-Strep in 5% Co2 at 37 ^ο^C. At 90% confluency spent media was aspirated off; cells were gently washed with PBS and incubated with DMEM only (no serum) for 24 to 72 hours to be conditioned with cell-secreted factors. At the end of incubation, conditioned media containing the breast cancer cell secretome was collected and passed through a 0.44 micron filter to remove any possible cell debris. Collected conditioned media was immediately flash frozen in liquid nitrogen, and stored in -80 ^ο^C.

For heat inactivation experiments, conditioned media was incubated at > 95 ^ο^C for 10 min and then allowed to cool at room temperature. Heat-inactivated conditioned was then mixed in a 1:1 ration with DMEM + 20%FBS + differentiation cocktail for in vitro adipogenesis experiments (see below).

Amicon ultra 100K centrifugal spin columns were used for size fractionation. Briefly conditioned media was transferred to a 100 kD Amicon spin column and centrifuged at 4000 g for 20 min. Filtrate and supernatant were collected separately and flash frozen in liquid nitrogen. For the in vitro adipogenesis experiments, the supernatant (top fraction) was reconstituted with DMEM to the initial volume of conditioned media loaded on spin column. Reconstituted supernatant and filtrate were separately mixed in a 1:1 ration with DMEM + 20% FBS + differentiation cocktail for in vitro adipogenesis experiments (see below).

#### *In vitro* Adipogenesis

3T3-L1 cells were grown to confluency in DMEM containing 10% Bovine Calf Serum, followed by another 2 days at confluency in DMEM containing 10% Bovine Calf Serum. Adipogenesis was then induced using DMEM containing 10% FBS and differentiation cocktail consisting of 1μg/ml insulin, 1μM Dex, and 0.5mM IBMX (differentiation media). After 2 days of differentiation cocktail, media was changed to DMEM containing 10% FBS and 1μg/ml insulin (maintenance media). Maintenance media was changed every 2-3 days for a total differentiation time of 4-8 days. Where indicated, DMEM containing the noted breast cancer cell secretome was used for the first 2 days of adipogenesis induction. Conditioned media was used at a one-to-one ratio with fresh DMEM, plus 10% FBS (final) and differentiation cocktail. To increase the dynamic range and better quantify rescue of adipogenesis using conditioned media from knockout cells, conditioned media collected from MDA-MB-468 sgRNA cells (and controls) was used at 100% (i.e., no fresh DMEM added) in the 3T3-L1 differentiation media. No conditioned media was used past the first 2 days of adipogenesis induction (i.e., in the maintenance media). Where indicated, differentiation media was supplemented with recombinant ZAG (R&D Systems #4764-ZA) or TSP-1 (R&D Systems #3074-TH). No conditioned media was used in the recombinant protein experiments.

Mouse primary preadipocytes were grown to confluency in DMEM containing 10% FBS. Adipogenesis was induced using DMEM containing 10% FBS and differentiation cocktail consisting of insulin, dexamethasone, and IBMX. After 3 days of differentiation cocktail, media was changed to DMEM containing 10% FBS and 1μg/ml insulin. Maintenance media was changed every 2-3 days. Breast cancer cell conditioned media was used during the first three days of adipogenesis induction where indicated (i.e., in differentiation media). TGFβ at 2ng/ml (Miltenyi Biotec, 130-095-067) treatment was used during the first three days of adipogenesis induction where indicated.

#### Isolation of primary preadipocytes

Mice were euthanized with isoflurane and secondary cervical dislocation. Inguinal white adipose tissue was dissected out and lymph nodes were removed. Tissue was removed and minced in HBBS (Gibco, 14025-92) and incubated in Collagenase Buffer (3,000 U/ml type II collagenase powder (Sigma, C6885), 100 U/ml DNase 1 (Roche, 10104159001), 1 mg/ml poloxamer 188 (P-188) (Sigma P5556), 1 mg/ml BSA, 20 mM HEPES buffer, and 1 mM CaCl_2_ in Medium 199 with Earle’s salts (Sigma, M4530)) for 15 min shaking (250 rpm) at 37LC. Equal volume of FBS was added to collagenase buffer to neutralize. Digested samples were strained through a sieve followed by centrifugation at 4LC (1300rpm for 10 minutes).

The cell pellet supernatant was resuspended in FACS buffer (PBS, 2% FBS, 1% PenStrep, 1% GlutaMAX) and strained through a 70 micron filter, followed by centrifugation at 4 LC (1300 rpm for 8 minutes). Pellet was resuspended in 37 LC Red Lysing Buffer (Sigma, R7767) for one minute, followed by inactivation using DMEM complete growth media (DMEM, 10% FBS, 1% PenStrep, 1% GlutaMAX). Cells were centrifuge at 4LC (1300rpm for 7 minutes) and resuspended into 5 ml of wash solution (300ug/ml DNase in sterilized water, 70% DPBS with MgCl_2_ and CaCl_2_ (Sigma, D8662)), followed by centrifugation at 4LC (1300rpm for 6 minutes). Cell pellet containing primary preadipocytes were washed with FACS buffer and resuspended into 200ul FACS Buffer.

For flow sorting ASPCS, antibodies were added to 200 ul of cell suspension in FACS buffer and incubated for 30 minutes on ice. Cells were then centrifuge at 4 LC (1300rpm for 5 minutes) and resuspended in 500 ul FACS Buffer. Cells were then sorted on BD FACS Aria Cell sorter. ASPCs (Lin-CD34+ Sca1+ CD29+) were resuspended in DMEM containing 10% FBS, 1% Pen/Strep, and 1% GlutaMAX. Antibodies used were the following: CD45 PE-Cy7 (eBioscience, 25-0451-82), CD31 PE-Cy7 (eBioscience, 25-0311-82), Ter119 PE-Cy7 (eBioscience, 25-5921-82), CD34 PE (BD Pharmingen, 551387), Sca1 APC (Biolegend 108112) and CD29 FITC (Invitrogen, 11-0291-82).

#### Live Cell Imaging Analysis for lipid accumulation and proliferation

Live imaging for kinetics and end-point quantification of adipogenesis was performed using the IncuCyte Live Cell Analysis Imaging System (Essen Bioscience) with images acquired every 4h using the 10x objective. Cells were differentiated as described above and supplemented with 2µM BODIPY 493/503 throughout the adipogenesis time course. Green fluorescence images were acquired every 4h using default IncuCyte setting. Total green fluorescence intensity was determined from a green fluorescent mask generated by the IncuCyte Zoom Analysis Software. For end-point analysis, final total green fluorescence intensity values (Day 4 or Day 5 of differentiation) were normalized to “no differentiation control” and “DMEM control differentiation” (no secretome).

Proliferation was assessed using a confluency mask generated by the IncuCyte Zoom Analysis Software using phase images.

#### Oil Red O staining and quantification

Cells are fixed in 4% PFA/PBS for 10min at room temperature, followed by 3 rinses in PBS. Samples were incubated in 60% isopropanol for 5min at room temperature and then allowed to dry completely. Samples are then incubated in freshly diluted 60% Oil Red O staining solution in water (stock is 0.5% Oil Red O (Sigma, 00625) in isopropanol) for 20min at room temperature, followed by 3 rinses in water. Samples were allowed to dry completely and imaged. To quantify, Oil Red O was extracted by incubating dried samples stained on the same day in 100% isopropanol for 5min at room temperature and absorbance was measured at 510nm.

#### Quantitative Real time PCR

RNA was extracted using the RNeasy Mini Kit (QIAGEN, 74104) and cDNA was synthesized using M-MLV Reverse Transcriptase (Invitrogen, 28025-013). Quantitative real time PCR was performed using TaqMan Probes (Invitrogen) and the TaqMan Gene Expression Master Mix (Applied Biosystems, 4369016) in 96-well MicroAmp Optical reaction plates (Applied Biosystems, N8010560).

#### Proteomics analysis

Digestion of in-gel proteins: Proteins were reduced with 5 mM DTT for 45 minutes at 60 °C and then alkylated with 10 mM IAA for 30 minutes at room temperature in the dark. Trypsin/LysC mixture was added to protein sample in a 1:100 ratio. Proteins were digested for overnight at 38 °C. The digestion was quenched by acidification with 1% formic acid to a pH of 2-3 and the peptides concentrated *en vacuo* to a final volume of 5 µL.

LC-MS/MS Analysis: Reversed-phase nano-LC-MS/MS was performed on an UltiMate 3000 RSLCnano system (Dionex) with a PharmaFluidics μPAC micro-chip based trapping column and a 50 cm equivalent PharmaFluidics μPAC micro-chip based column (Pharmafluidics, Ghent, Belgium). Peptide fractions were reconstituted in 100 μL of 0.1% formic acid in water. Five µL of the samples were injected into the liquid chromatograph. A gradient of reversed-phase buffers (Buffer A: 0.1% formic acid in water; Buffer B: 0.1% formic acid in acetonitrile) at a flow rate of 0.5 mL/min was set-up. The LC run lasted for 90 minutes with a starting concentration of 1% buffer B increasing to 28% over the initial 72 minutes with a further increase in concentration to 50% over 18 minutes. A final ramp up to 95% took place over 5 minutes and was held for 10 minutes before ramping down to 1% B over 2 minutes and re-equilibrating for 10 minutes. A

QExactive HF (ThermoFisher Scientific) coupled to a Flex nano spray source was employed with the following settings for MS1; resolution 120,000, AGC target 1e5, maximum IT 50 ms, scan range 375-1400 m/z. MS2 settings were; resolution 60,000, AGC target 1e5, maximum IT 100 ms, isolation window 1.2 m/z. Top 15 DDA analysis was performed with NCE set to 32.

Analysis of MS/MS Data: Proteome Discoverer (Version 2.4) was used for database searching and protein identification. For these samples the Uniprot database was searched with Homo sapien taxonomy selected. The parameters used for the Mascot searches were: no enzyme digest; two missed cleavages; carbamidomethylation of cysteine set as fixed modification; oxidation of methionine, acetylation of n-terminus, amidation of c-terminus, and deamidated of NQ, were set as variable modifications; and the maximum allowed mass deviation was set at 11 ppm.

#### Sample preparation and immunoblot

Cells were lysed on ice in RIPA buffer (Thermo Scientific, J62524) containing protease inhibitor (Thermo Scientific, Pierce A32965) and phosphatase inhibitor (Thermo Scientific, Pierce A32957). Lysed cells were spun down and supernatant was incubated with NuPAGE™ LDS sample buffer (Invitrogen, NP0007) for 5 minutes at 95 LC. Proteins were separated using NuPAGE™ 4 to 12%, Bis-Tris, 1.0–1.5 mm, Mini Protein Gels (Thermo Fisher Scientific, NP0321BOX) in NuPAGE™ MES SDS Running Buffer (Thermo Fisher Scientific NP0002), followed by transfer onto nitrocellulose membranes (BIO RAD, 1620115) using Towbin Buffer (2.5 mM Tris, 19.2 mM glycine, pH 8.3) containing 20% methanol. Membranes were blocked with blocking buffer (3% milk, TBST buffer (20 mM Tris, 150 mM NaCl, 0.1% Tween 20, pH 7.5)) for 30 minutes at room temperature, followed by incubation with primary antibody in 1% milk and in TBST buffer overnight (16 hour) at 4 LC. Membrane was washed 3 times in TBST buffer for 10 minutes and incubated with secondary IRDye antibodies (1:5000 LI-COR) in blocking buffer (1% milk, TBST, 0.001% SDS) for 30 minutes at room temperature. Membranes were washed 3 times in TBST buffer and scanned on an Odyssey CLx Imaging System (LI-COR). The following antibodies were used: ZAG (Santa Cruz, sc-13585 1:200), P-Histone H3 (Cell signaling, 34655 1:1000) IRDye 680 RD (LiCor, 926-68072 1:2000).

#### Clinical data analysis of human breast cancer patients

We used cBioPortal (Cerami et al., 2012; de Bruijn et al., 2023; Gao et al., 2013) to analyze the two breast cancer patient gene expression data sets TCGA (Cancer Genome Atlas, 2012) and Metabric (Curtis et al., 2012; Pereira et al., 2016; Rueda et al., 2019). These data sets were chosen because of availability of gene expression data as well as ER, PR, and HER2 status for all patients. All analyses were performed for both data sets combined or individually, groups were classified as all breast cancer, ER-positive breast cancer (ER Status positive), or TNBC (ER Status negative, HER2 Status negative, PR Status negative). For ZAG expression, mRNA expression z-scores relative to all samples (log microarray) were plotted for each group. For clinical outcome, ZAG/AZGP1 expression (mRNA expression z-scores relative to all samples (log microarray) was binned by quartiles, and survival was plotted. Analysis was repeated for each group (all breast cancer, ER-positive breast cancer, TNBC) and the two data sets combined and individually.

#### ELISA

ELISA for ZAG was performed according to DuoSet ELISA development system catalog no. DY4764 instructions. Briefly wells were coated with 100 ul diluted capture antibody, sealed and incubated overnight at room temperature (RT). Next day, washing steps were performed as per instructions, and wells were blocked using supplied blocking buffer for 1 hour at RT. Washing steps were repeated, and wells were treated with reagent diluent and incubated for 2 hours. Wells were washed and detection antibody was added to wells, which were covered and incubated at RT for 2hr. Post-incubation, washes were performed, and working dilution of Streptavidin-HRP was added to wells, which were covered and incubated for 20min at RT, avoiding light exposure. Washes were performed, followed by addition of substrate solution and a 20 min incubation. Stop solution was added to terminate the reaction, followed by thorough mixing by tapping and measuring OD at 450 nm in a microplate reader (Biotek Neo2).

#### Xenograft studies

Orthotopic xenograft studies were performed in 5 weeks old NRG mice. Two independent studies were done weeks apart, and each experimental cohort included total of 10 mice (5 for control cells + 5 for ZAG depleted cells). 5 x10^6 cells of MDA-MB-468 sgSafe (control) or MDA-MB-468 sgZAG were injected with Matrigel into the 4^th^ mammary fat pad of female NRG mice. During the course of the study, mice were weighed 1X per week, and tumors were measured in millimeters by length and width 2x per week to calculate the tumor volumes. At the end of the study, all mice from each of the experimental cohorts were euthanized together, and tumors with surrounding adipose tissue were collected, embedded, and stained for analysis.

#### Immunofluorescence staining

Tumors and surrounding adipose tissue were paraffin embedded and sectioned by the ARUP Histology Core Laboratory. For deparaffinization of tumor sections, slides were warmed at 70 LC for 45 minutes. The slides were then incubated in the following series of solutions at room temperature: 100% xylene (Sigma-Aldrich, 534056) for 15 minutes (two times), 75% xylene and 25% ethanol (Decon Labs, 64-17-5) for 7 minutes, 50% xylene and 50% ethanol for 5 minutes, 25% xylene and 75% ethanol for 5 minutes, 100% ethanol for 5 minutes (two times), 75% ethanol for 5 minutes, 50% ethanol for 5 minutes, 25% ethanol for 5 minutes, then tumor section slides were washed in 100% water for 5 minutes. Heat-induced epitope retrieval was used for the following antibodies: Ki67 (Cell Signaling D35B, 1:200), Alpha-Smooth Muscle Actin (Invitrogen, 14-9760-82, 1:200); briefly, 10mM sodium citrate buffer (10 mM Sodium citrate, 0,05% tween 20, Ph 6.0) was warmed to 95 LC. Slides were then incubated in sodium citrate buffer for 15 minutes, followed by cooling at room temperature for 20 minutes.

For immunofluorescence staining, slides were washed 3 times for 5 minutes with PBS at room temperature. Slides were then blocked for 60 minutes with blocking buffer (1% PBSA (1% BSA (Sigma-Aldrich, A7906) in PBS)) for 60 minutes. Slides were then incubated in 1% PBSA with primary antibody for 60 minutes, followed by washing the sections in 1% PBSA 3 times for 10 minutes at room temperature. Slides were incubated with secondary antibody (Alexa Fluoro, 1:200 dilution) in 1% PBSA for one hour. Slides were then incubated DAPI (1:1000) for 5 minutes, followed by washing the sections with 1% PBSA 3 times for 10 minutes. Fluoromount - G (Southern Biotech, 0100-01) was used to mount slides with cover slips. Primary antibodies used were the following: Ki67 (Cell Signaling D35B, 1:200), Alpha-Smooth Muscle Actin (Invitrogen, 14-9760-82, 1:200).

#### Sirius red staining

Slides with sections of tumors and surrounding adipose tissue were deparaffinized and hydrated in distilled water as described above. Slides were stained with Picro Sirius Red Staining Kit as per manufacturer’s instructions (Abcam, ab150681). Slides were mounted with cover slides using Cytoseal XYL (Espredia, 8312-4).

#### Imaging

Slides were imaged on a Zeiss Axioscan Z1 and analyzed using NIS-Elements (NIKON) software. To quantify fibrosis, a threshold intensity for Sirius Red staining was set and the Sirius Red positive area was determined in the region of interest of adipose tissue (within 2mm surrounding the tumor). Percent positive Sirius Red was calculated per tumor/adipose tissue sample and averaged. To quantify proliferation, a threshold intensity was set for Ki67 immunofluorescence stain. Percent positive Ki67 positive nuclei was calculated per tumor samples and averaged. To quantify fibrosis using alpha SMA staining, a threshold intensity was set and alpha SMA positive area was determined within the adipose tissue region of interest (within 2mm of the tumor). Percent positive alpha SMA area was calculated per tumor/adipose tissue and averaged. Adipocyte size was quantified on hematoxylin and eosin (H&E)-stained slides using the Adiposoft (Galarraga et al., 2012) plugin on ImageJ (NIH). Briefly, regions of interest were selected based on ROIs were chosen (minimum diameter of 20 pixels, max diameter of 200 pixels; pixel size 0.878). Frequency distribution of adipocyte diameter was determined per tumor/adipose tissue and averaged.

### QUANTIFICATION AND STATISTICAL ANALYSIS

Statistical parameters including the statistical test used, exact value of n, what n represents, and the distribution and deviation are reported in the figures and corresponding figure legends. Most data are represented as the mean ± standard deviation, data points show independent biological replicates, and the p-value was determined using unpaired two-tailed Student’s t tests, one-way ANOVA, or two-way ANOVA.

Unless otherwise stated, statistical analyses were performed in GraphPad Prism.

