## Supplemental Figures for "Zinc Alpha-2-Glycoprotein (ZAG/AZGP1) secreted by triple-negative breast cancer promotes tumor microenvironment fibrosis"

A

| Luminal A | Luminal B | Her2+ | Triple Negative |
|---|---|---|---|
| MCF7<br>T47D | BT474<br>UACC812 | SKBR3<br>MDA-MD-453 | MDA-MB-468<br>BT-20<br>MDA-MB-231<br>Hs578T |

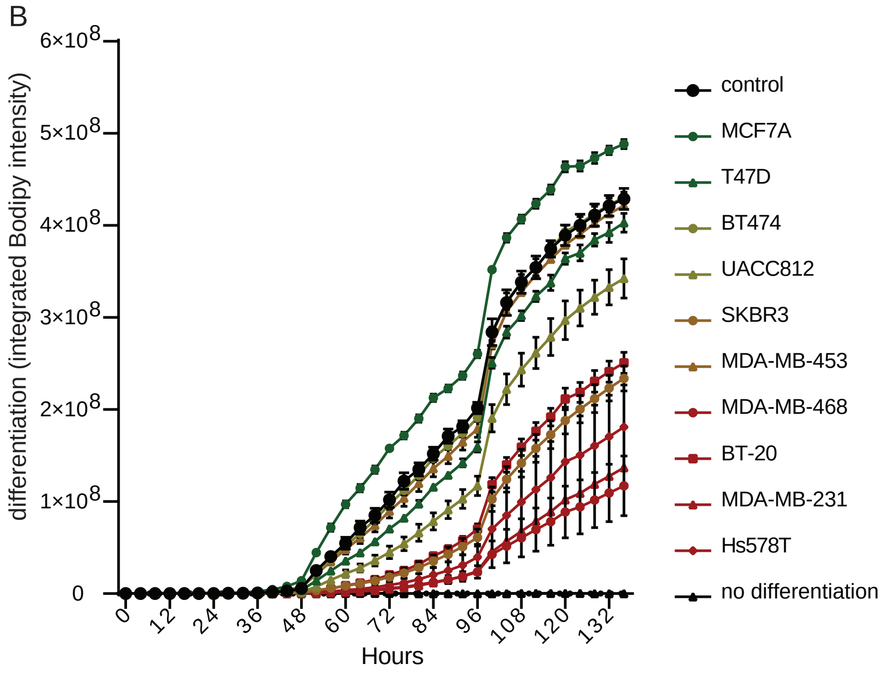


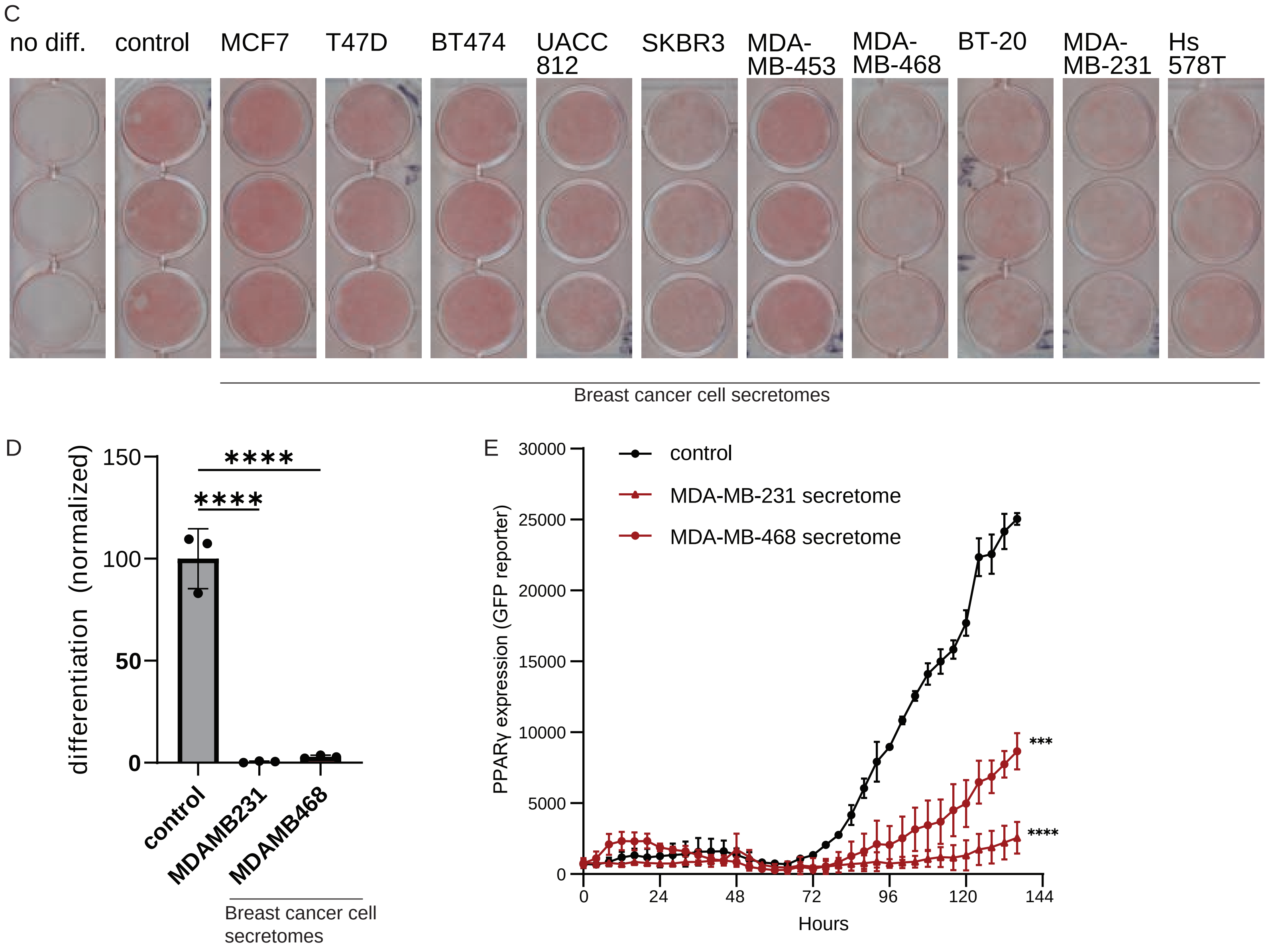


Figure S1

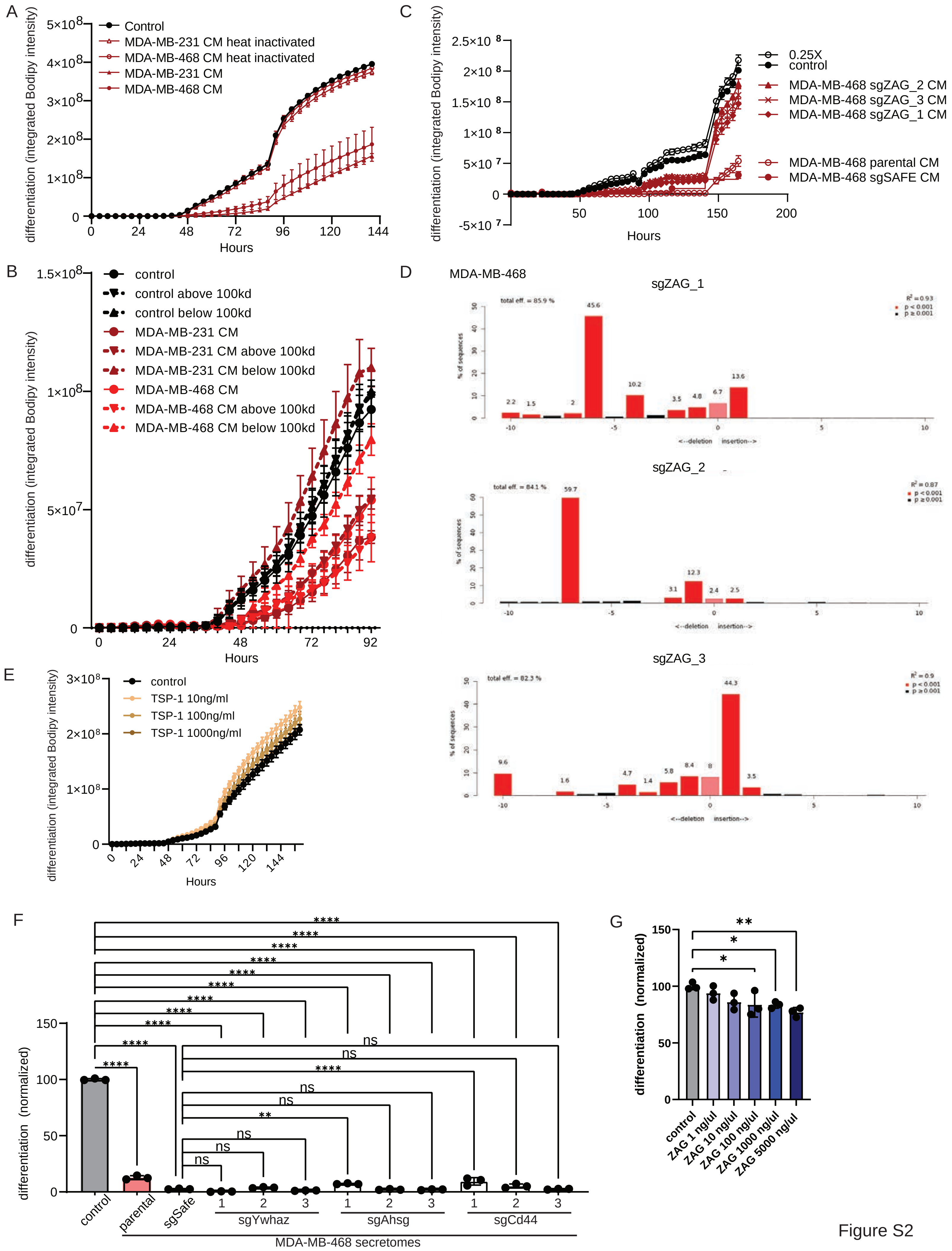
A
differentiation (integrated Bodipy intensity)
Hours
Control
MDA-MB-231 CM heat inactivated
MDA-MB-468 CM heat inactivated
MDA-MB-231 CM
MDA-MB-468 CM
B
differentiation (integrated Bodipy intensity)
Hours
control
control above 100kd
control below 100kd
MDA-MB-231 CM
MDA-MB-231 CM above 100kd
MDA-MB-231 CM below 100kd
MDA-MB-468 CM
MDA-MB-468 CM above 100kd
MDA-MB-468 CM below 100kd
C
differentiation (integrated Bodipy intensity)
Hours
0.25X
control
MDA-MB-468 sgZAG_2 CM
MDA-MB-468 sgZAG_3 CM
MDA-MB-468 sgZAG_1 CM
MDA-MB-468 parental CM
MDA-MB-468 sgSAFE CM
D
MDA-MB-468
sgZAG_1
total eff. = 85.9 %
sgZAG_2
total eff. = 84.1 %
sgZAG_3
total eff. = 82.3 %
% of sequences
<--deletion insertion-->
E
differentiation (integrated Bodipy intensity)
Hours
control
TSP-1 10ng/ml
TSP-1 100ng/ml
TSP-1 1000ng/ml
F
differentiation (normalized)
control
parental
sgSafe
sgYwhaz
sgAhsg
sgCd44
MDA-MB-468 secretomes
G
differentiation (normalized)
control
ZAG 1 ng/ul
ZAG 10 ng/ul
ZAG 100 ng/ul
ZAG 1000 ng/ul
ZAG 5000 ng/ul

Figure S2

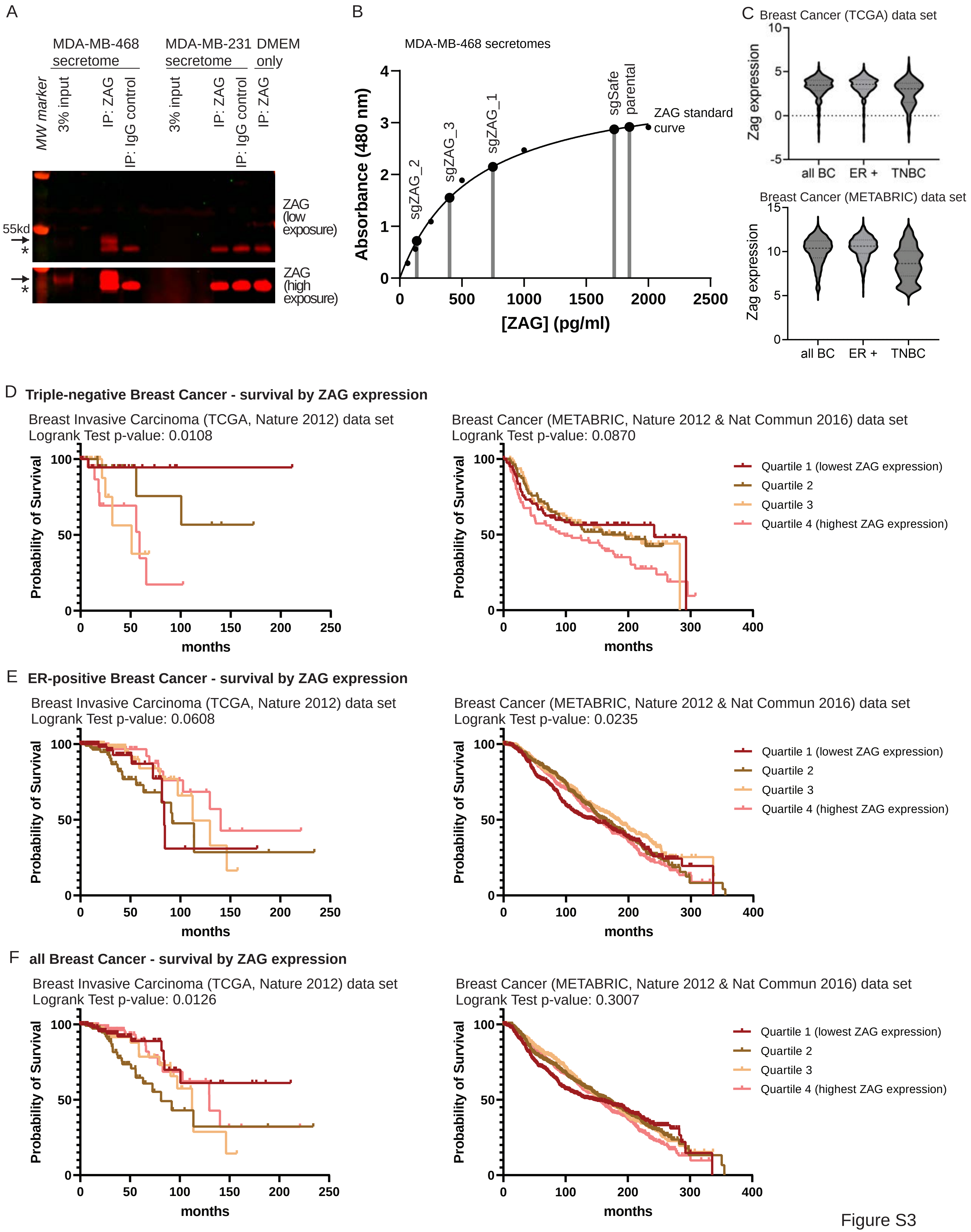
A
MDA-MB-468 secretome
MDA-MB-231 secretome
DMEM only
MW marker
3% input
IP: ZAG
IP: IgG control
3% input
IP: ZAG
IP: IgG control
IP: ZAG
55kd
*
ZAG (low exposure)
ZAG (high exposure)
B
MDA-MB-468 secretomes
Absorbance (480 nm)
[ZAG] (pg/ml)
sgZAG_2
sgZAG_3
sgZAG_1
sgSafe
parental
ZAG standard curve
C
Breast Cancer (TCGA) data set
Zag expression
all BC
ER +
TNBC
Breast Cancer (METABRIC) data set
Zag expression
all BC
ER +
TNBC
D
Triple-negative Breast Cancer - survival by ZAG expression
Breast Invasive Carcinoma (TCGA, Nature 2012) data set
Logrank Test p-value: 0.0108
Probability of Survival
months
Breast Cancer (METABRIC, Nature 2012 & Nat Commun 2016) data set
Logrank Test p-value: 0.0870
Probability of Survival
months
Quartile 1 (lowest ZAG expression)
Quartile 2
Quartile 3
Quartile 4 (highest ZAG expression)
E
ER-positive Breast Cancer - survival by ZAG expression
Breast Invasive Carcinoma (TCGA, Nature 2012) data set
Logrank Test p-value: 0.0608
Probability of Survival
months
Breast Cancer (METABRIC, Nature 2012 & Nat Commun 2016) data set
Logrank Test p-value: 0.0235
Probability of Survival
months
Quartile 1 (lowest ZAG expression)
Quartile 2
Quartile 3
Quartile 4 (highest ZAG expression)
F
all Breast Cancer - survival by ZAG expression
Breast Invasive Carcinoma (TCGA, Nature 2012) data set
Logrank Test p-value: 0.0126
Probability of Survival
months
Breast Cancer (METABRIC, Nature 2012 & Nat Commun 2016) data set
Logrank Test p-value: 0.3007
Probability of Survival
months
Quartile 1 (lowest ZAG expression)
Quartile 2
Quartile 3
Quartile 4 (highest ZAG expression)

Figure S3

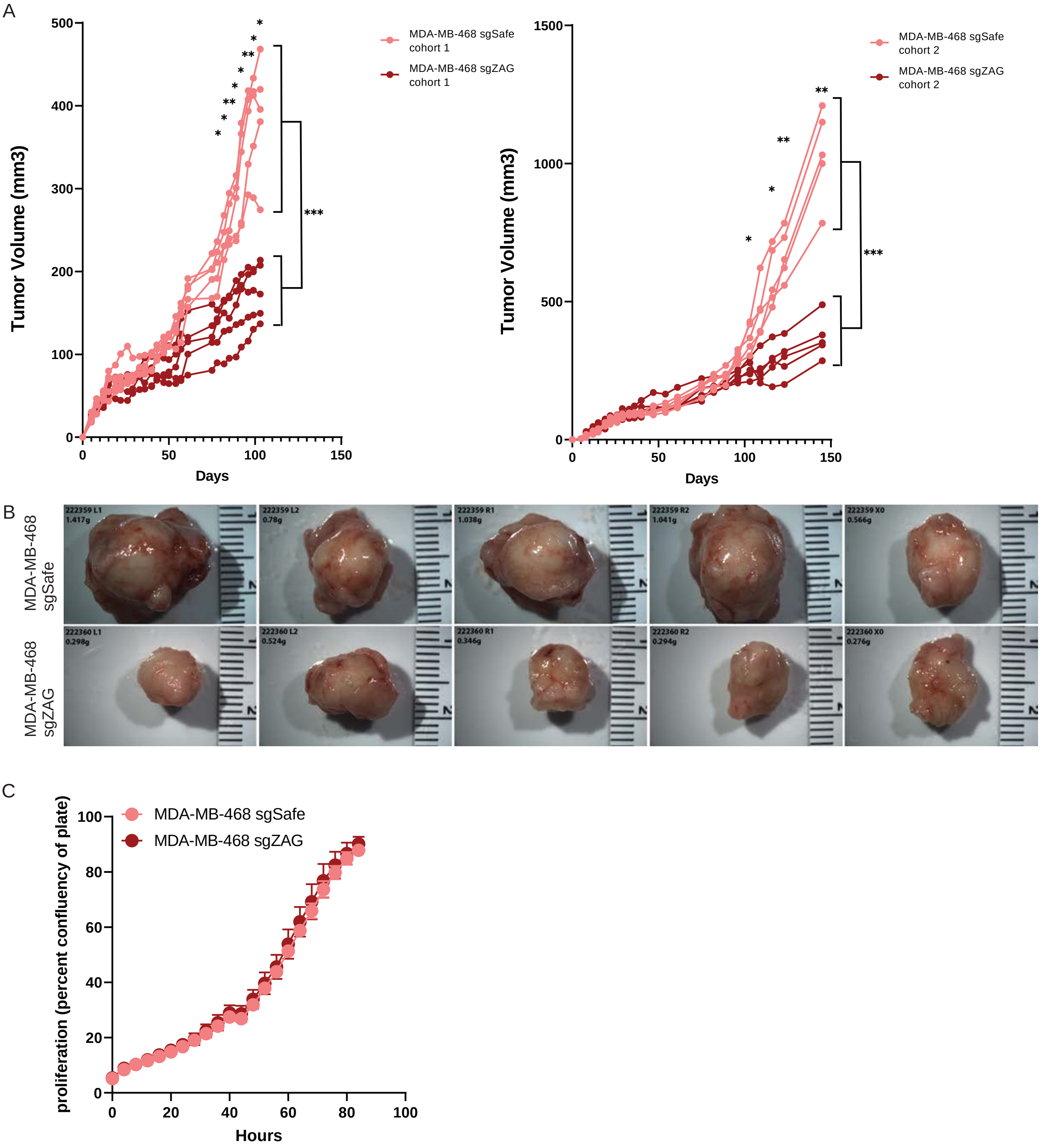
A
Tumor Volume (mm3)
Days
MDA-MB-468 sgSafe cohort 1
MDA-MB-468 sgZAG cohort 1
***
MDA-MB-468 sgSafe cohort 2
MDA-MB-468 sgZAG cohort 2
***
B
MDA-MB-468 sgSafe
MDA-MB-468 sgZAG
222359 L1 1.417g
222359 L2 0.78g
222359 R1 1.038g
222359 R2 1.041g
222359 X0 0.566g
222360 L1 0.298g
222360 L2 0.524g
222360 R1 0.346g
222360 R2 0.294g
222360 X0 0.276g
C
proliferation (percent confluency of plate)
Hours
MDA-MB-468 sgSafe
MDA-MB-468 sgZAG

Figure S4

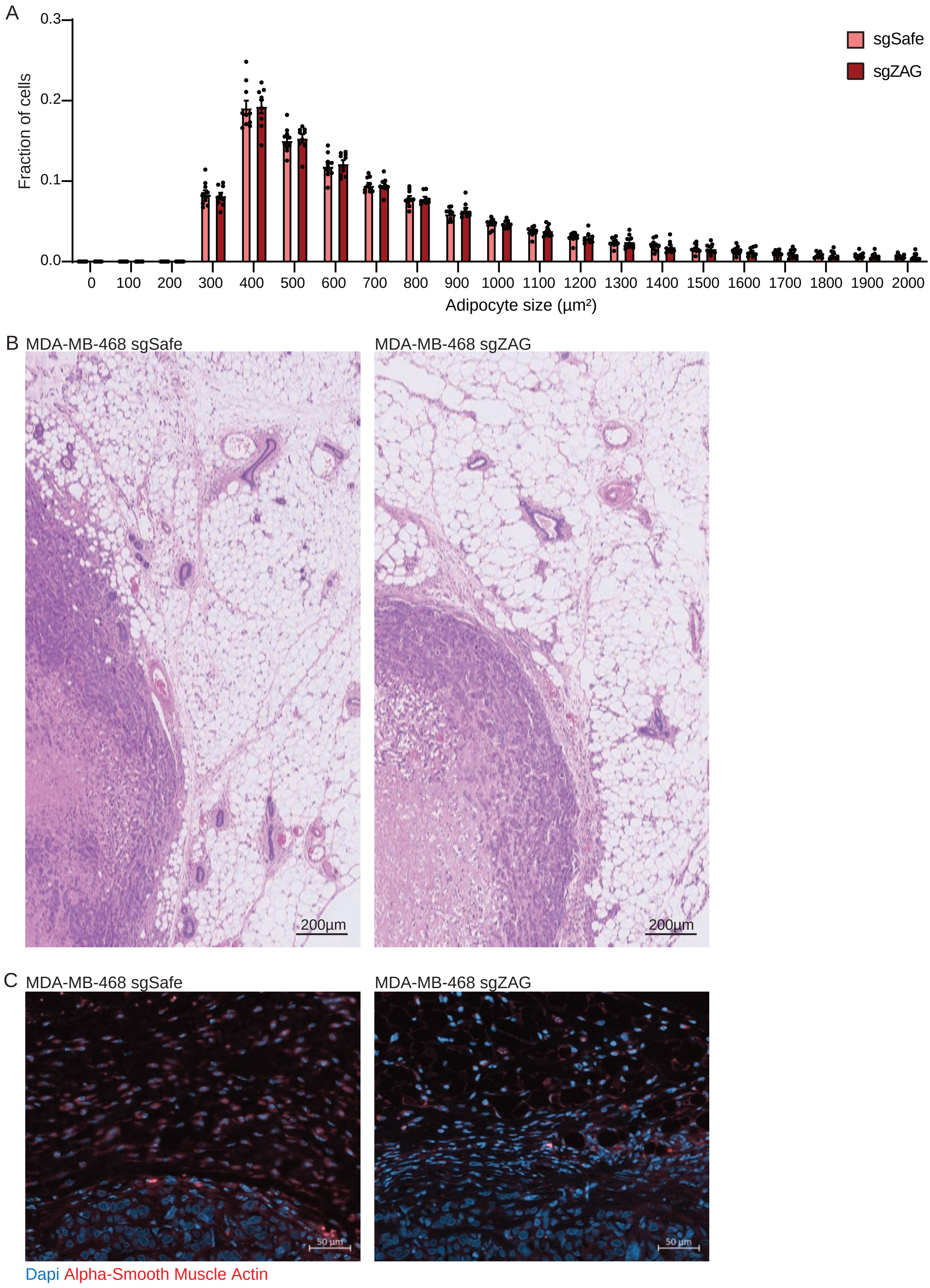
A
0.3
0.2
0.1
0.0
Fraction of cells
sgSafe
sgZAG
0
100
200
300
400
500
600
700
800
900
1000
1100
1200
1300
1400
1500
1600
1700
1800
1900
2000
Adipocyte size (µm²)
B MDA-MB-468 sgSafe
MDA-MB-468 sgZAG
200µm
200µm
C MDA-MB-468 sgSafe
MDA-MB-468 sgZAG
50 µm
50 µm
Dapi Alpha-Smooth Muscle Actin

Figure 5
